# Wet Adhesive Hydrogels to Correct Malacic Trachea (Tracheomalacia): A Proof of Concept

**DOI:** 10.1101/2022.11.29.518329

**Authors:** Ece Uslu, Vijay Kumar Rana, Sokratis Anagnostopoulos, Peyman Karami, Alessandra Bergadano, Cecile Courbon, Francois Gorostidi, Kishore Sandu, Nikolaos Stergiopulos, Dominique P. Pioletti

**Affiliations:** Laboratory of Biomechanical Orthopedics, Institute of Bioengineering, School of Engineering, EPFL, Lausanne, Switzerland; Laboratory of Hemodynamics and Cardiovascular Technology, Institute of Bioengineering, School of Engineering, EPFL, Lausanne, Switzerland; Department of Otorhinolaryngology, Airway sector, University Hospital, CHUV, Lausanne, Switzerland; Veterinary College, Bern, Switzerland; Department of Anesthesiology, University Hospital, CHUV, Lausanne, Switzerland

## Abstract

Tracheomalacia (TM) is a condition in which the anterior part of the trachea consisting of cartilage and/or the posterior part consisting muscle are too soft to ensure its mechanical support. This situation may result in an excessive and potentially lethal collapse of the airway in the newborns. Current treatment techniques include tracheal reconstruction, tracheoplasty, endo- and extraluminal stents, but are all facing important limitations.

To reduce the shortcomings of actual TM treatments, this work proposes a new strategy by wrapping an adhesive hydrogel patch extraluminally around a malacic trachea. To validate this approach, first a numerical model revealed that a hydrogel patch with sufficient mechanical and adhesion strength can potentially preserve the trachea’s physiological shape. Accordingly, a new hydrogel formulation was synthesized employing the hydroxyethyl acrylamide (HEAam) and polyethylene glycol methacrylate (PEGDMA) as main polymer network and crosslinker, respectively. These hydrogels provide excellent adhesion on wet tracheal surfaces, thanks to a two-step photo-polymerization approach. Ex vivo experiments revealed that the developed adhesive hydrogel patches can restrain the collapsing of malacic trachea under applied negative pressure. This study, to be confirmed by in vivo studies, is open to the possibility of a new treatment in the difficult clinical situation of tracheomalacia in newborns.

## Introduction

Tracheomalacia (TM) is characterized by excessive luminal collapse due to immature cartilage formation and floppiness of trachealis muscle^[1]^. It can either be congenital or acquired. Congenital TM is the most common tracheal abnormality seen in neonates at an incidence rate of 1:2100^[2]^. Whereas an acquired TM can emerge due to infection, inflammation, tracheostomy, trauma, and compression exerted by abnormal cardiovascular structures^[2]^. Essentially a healthy trachea contains 18-22 anterior C-shaped cartilage rings as well as posterior membranous part. The trachealis muscle in the membranous trachea moves toward cartilage rings during exhalation and forceful efforts (crying, coughing, laughing, *etc.*) in intrathoracic segment of the trachea, which helps the airflow and mucus clearance^[2]^. During such movements, the openings of the airway lumen are narrowed down by 10-20% as depicted in Fig. 1. Contrarily, if suffering from TM, the same cartilage rings tend to be U or bowed-shaped with broader and more dynamic trachealis muscles^[3]^, as portrayed in Fig. 1. These changes in tracheal shape are accountable for airway collapse during expiration and forceful efforts in an intrathoracic segment of the trachea. This condition prevents normal breathing and could be life-threatening, especially for newborns^[1]^. In neonates, infants, and small children, tracheal lumen obstruction up to 50% due to a posterior to anterior collapse (tracheal muscle movement towards the cartilage ring) can be physiological, collapse exceeding 50% obstruction will cause respiratory symptoms and collapse >75% will be critical. Clinically, the collapse in the trachea can be classified as mild, moderate, and severe if it shrinks by 25-50%, 50-75%, and >75%, respectively^[4,5]^.

**Figure 1:**
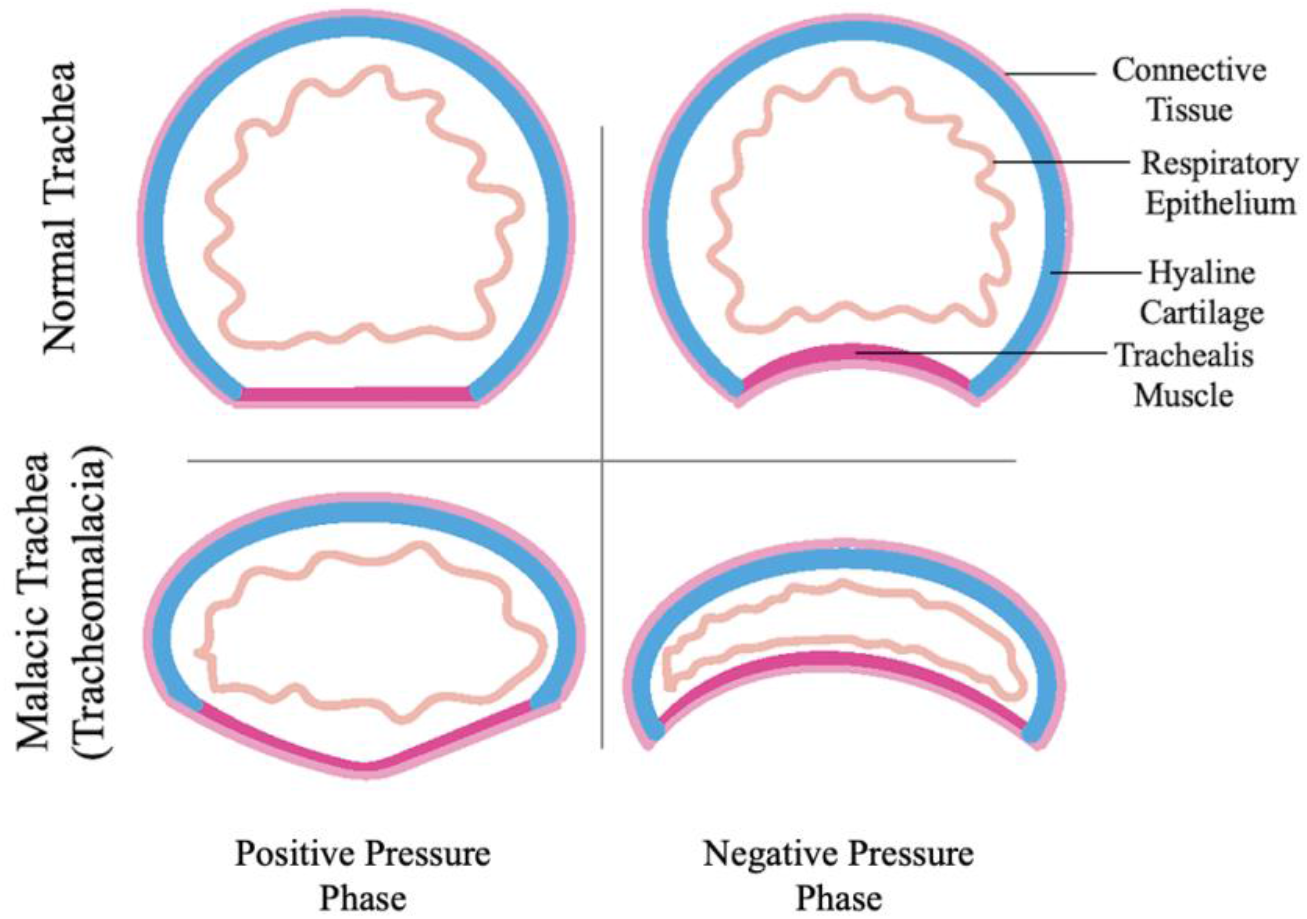
Representation of normal and malacic trachea during positive and negative pressure phase of the respiratory cycle.

The main clinical approach to solve TM is to correct the geometry of the malacic trachea and to prevent the airway collapse. Depending on the severity, surgical interventions and stenting are commonly used treatment methods for TM. Surgical interventions like, aorto- and sterno-pexy alleviate the pressure by placing static suspension sutures on the trachea and attempt to increase the airway area.

However, these techniques are time-consuming and may fail if the pexy sutures breakdown. Additionally, they are not indicated for TM cases requiring mechanical support as in a circumferential floppy airway, in form of external airway splints and endoluminal stents^[2,6–8]^. Alternatively, tracheal resection-anastomosis and slide tracheoplasty are used only for short segment TM which is seen seldom in clinics^[9]^. On the other hand, intraluminal stents (silicone, metallic and bioresorbable) and extraluminal splints (bioresorbable plates and 3D-printed splints) have also been considered to correct the malacic trachea shape and stabilize the airway collapse by supporting the trachea mechanically^[10]^. However, both techniques come with numerous challenges to encounter. Intraluminal stents lead to severe complications in children, including stent obstruction due to secretions and granulation tissue, stent-induced stenosis, migration and erosion^[10]^. Extraluminal splints are fixed outside the tracheal lumen using sutures and have the advantages of avoiding complications of the endoluminal stent and support the structure externally ^[11]^.

However, the splint fixing sutures can cause obstructing granulations at the contact point inside the trachea and their rigidity could prevent the natural and dynamic neck movements in the child. To understand TM condition broadly, we can look into a ubiquitous engineering problem in the piping industry. Pipes that are being used in the oil and gas industries often tumble if excess external pressure is applied^[12]^. The primary cause of pipe collapsing is inadequate mechanical properties of the materials utilized to manufacture them^[13,14]^ and to prevent such unforeseen collapses, materials with better mechanical strengths could be paramount. Alternatively, depending upon the severity of damage, an internal or external mechanical support to optimally support the pipe collapse could be a solution.

Inspired by the experiences developed in the pipe industry, here, we propose a proof of a new conceptual approach to correct TM where a malacic trachea can be supported mechanically by wrapping an adhesive hydrogel patch extraluminally. Hydrogels are 3D water-swollen polymers network that mimic the properties of many tissues^[15]^. These viscoelastic biocompatible materials allow the diffusion of oxygen, nutrients, and other molecules, which is an ideal environment for the cells to grow^[16,17]^. We envisioned that adhesive hydrogel could be employed to mitigate many surgical issues while treating TM. For example, adhesion can avoid placement of extraluminal stent fixation sutures, reduce the associated trauma to the surrounding tissues, and the softness of the hydrogel can avoid unnecessary pain during natural movement of the trachea in newborns compared to stiff materials.

Herein, first, a numerical model has been developed to test the hypothesis that airway collapse can be prevented by providing external mechanical support to the trachea with an adhesive hydrogel patch. The model showed that the application of an adhesive hydrogel patch has the potential to preserve a more physiological shape of the trachea by constraining the posterior tracheal membrane folding. Accordingly, a new adhesive hydrogel is formulated employing hydroxyethyl acrylamide (HEAam) and polyethylene glycol methacrylate (PEGDMA) as the main polymer network and crosslinker, respectively. These hydrogels provided robust adhesion on wet tracheal surface owing to a two-step polymerization approach that help to anchor the surface much better. To evaluate the potential of the new approach in the treatment of TM, *ex vivo* experiments were performed.

## Results and Discussion

A numerical model based on finite element analysis (FEA) was developed to reinforce the hypothesis that local external mechanical support can hinder the airway collapse, and to understand the deformation and stress response of the trachea under maximum negative pressure with and without wrapping a hydrogel patch. A simple geometry was obtained from μ-CT scans of the rabbit tracheas harvested in slaughterhouses and modelled on AutoCAD Inventor. Thereupon, the trachea was placed inside the μ-CT machine, and a syringe-based vacuum system with an electronic manometer was connected to apply pressure (+/-) to deform it for later use, as shown in Fig. 2a. To accurately capture the geometrical features of the trachea (the radially variable thickness, the asymmetric shape, *etc.*), five equidistant scans were used as a guideline to sketch the exact profile at the corresponding cross-sections (Fig. 2b-c). A 3D model was then fitted through these sketches using a loft function, as shown in Fig. 2d. Details are given in the methods section. Due to their fibrous composition, it has been previously assumed in the literature that the tracheal components are incompressible, isotropic, and possess hyper-elastic properties^[18]^. The most accurate model is the Mooney-Rivlin model, which has been employed to represent the hyper-elastic properties of all three tracheal components (cartilage, muscle, connective tissue)^[19,20]^. Therefore, it has also been selected for this study. The strain-energy function generalized model can be defined as:

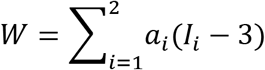

**Figure 2:**
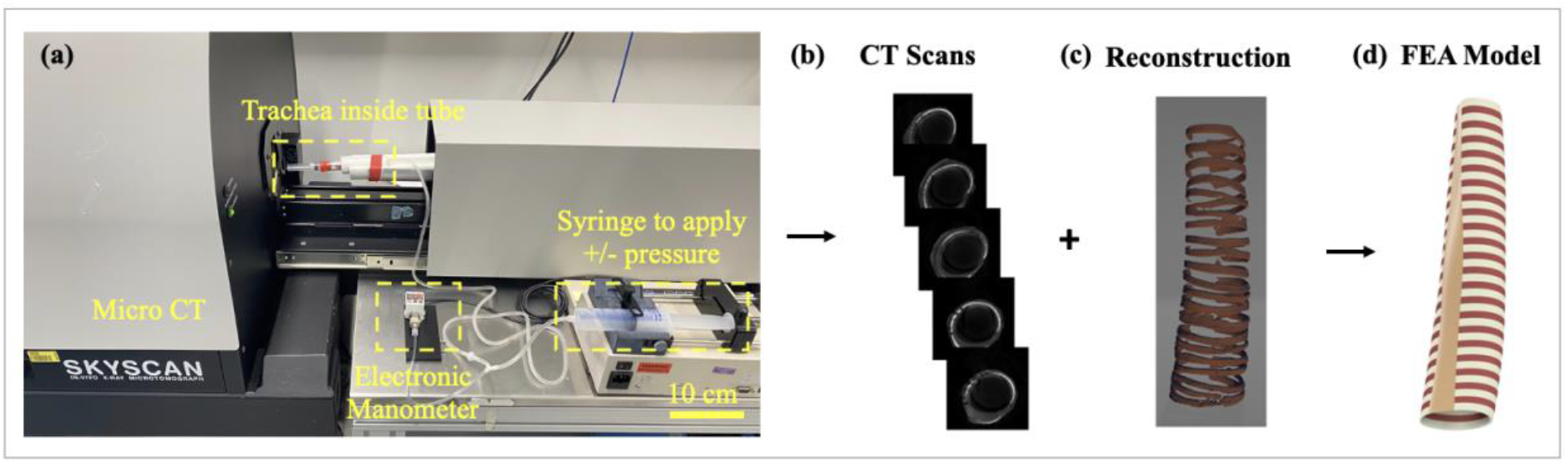
**(a)** Micro-CT (μ-CT) and a set-up to fix the rabbit trachea inside the machine and create the negative pressure using a syringe filled with air, **(b-d)** a 3D CAD design process of the trachea: **(b)** Five equidistant crosssections obtained from the μ-CT scanner, **(c)** 3D reconstruction of the cartilage-to-tissue volume ratio, **(d)** generation of the final realistic model of equivalent geometrical properties.

Where *W* is the strain energy, *a_i_* are the constants of the material and *I_i_* are the strain invariants. Additional information on the implementation of the Mooney-Rivlin model in Ansys can be found in the experimental and methods section.

To achieve mesh-independent results, a parametric study of multiple refined steps was conducted. The maximum deformation and von Mises stress were converged after about 2·x10^5^ quaddominant elements. The indicative results of deformation and von Mises stresses under −15 mmHg pressure are shown in Fig. 3(a-d). The maximum deformation is experienced by the tracheal membrane, which is primarily responsible for the air blockage in the malacic tracheas. Expectedly, we noticed that the tracheal cartilage rings carry maximum stresses during tracheal deformation (or respiration).

**Figure 3:**
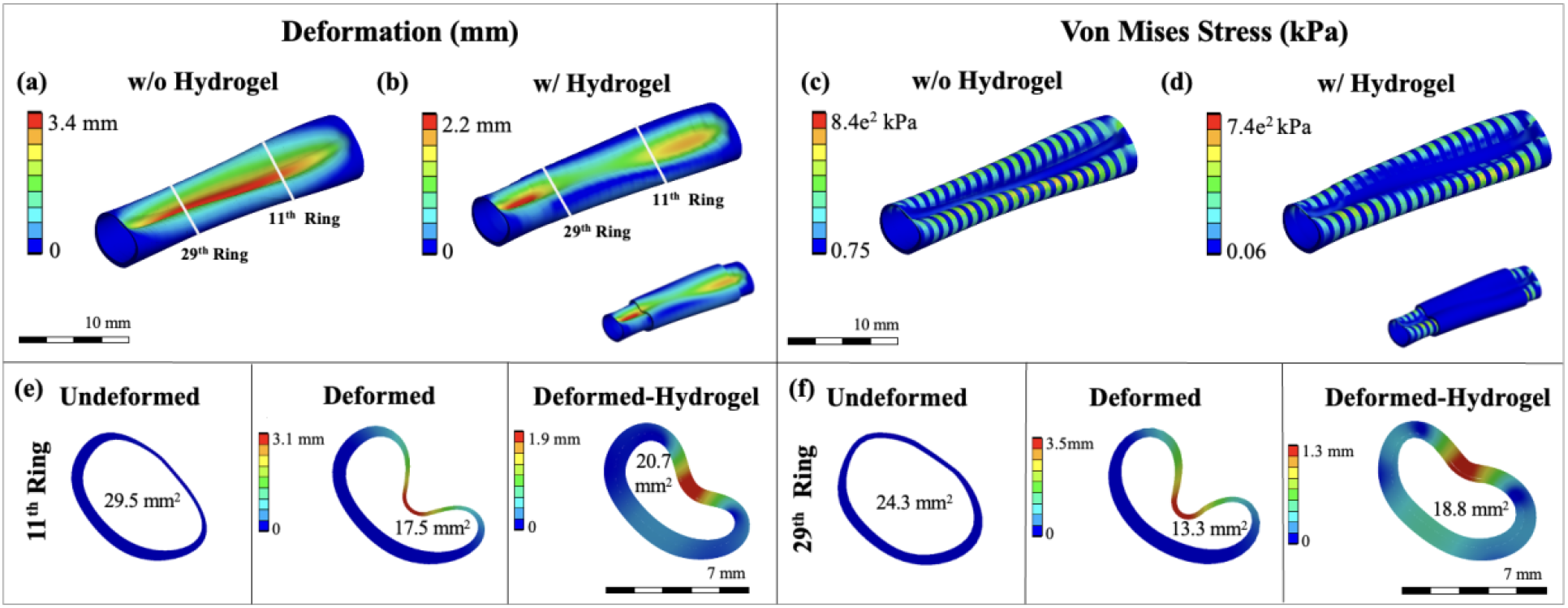
Indicative results of the total deformation of a trachea **(a)** without and **(b)** with hydrogel patch, and the stress distribution **(c)** without and **(d)** with hydrogel patch along with the 3D trachea model at −15 mmHg. Results show that with the presence of the hydrogel total deformation and stress decreases along the trachea. **(e-f)** Cross-sections of the rabbit trachea model at the 11^th^ and 29^th^ tracheal rings and subjected to −15 mmHg pressure. Numerical analysis indicate that applying a hydrogel patch to this tracheomalacic model inhibits the posterior collapsing.

We further simulated a trachea after wrapping an adhesive hydrogel patch to it under identical conditions. The hydrogel patch has been modeled as a homogeneous material with Young’s modulus (E) equal to 0.07 MPa, and a thickness of 0.7 mm (equal to the maximum thickness of the trachea ring). The membrane folding along with the suction of the thinner cartilage endings have been captured by the large deformation model. As shown in Fig. 3b and d, the deformation and stress values were reduced in a hydrogel-wrapped trachea. This strongly suggests that the hydrogel patch has the potential to constrain membrane folding. Specifically, the qualitative and quantitative results of the model show the deformation in the 11th and 29th tracheal rings under loading, as depicted in fig. 3e-f. The open area of the undeformed rings was 29.5 and 24.3 mm^2^ and decreased to 17.5 and 13.3 mm^2^ under −15 mmHg pressure, respectively. Interestingly, the open airway area (degree of collapse) was improved by ~11% and ~23% once a hydrogel patch is applied to the11th and 29th tracheal rings, respectively.

Based on the numerical findings, adhesive hydrogels were synthesized after photo-polymerizing the mixture of hydroxyethyl acrylamide (HEAam) monomers and cross-linkers (Polyethylene glycol dimethacrylate (PEGDM) or Gelatin methacrylate GelMA) at specific ratios using lithium phenyl-2,4,6-trimethylbenzoylphosphinate (LAP) as photoinitiator. The precursor solution was poured in a mold (15×15×0.7 mm^3^) and illuminated under 405 nm light for 2 minutes to afford a covalently crosslinked 3D network of adhesive hydrogel patch, as shown in Fig. 4a. A second polymerization was performed after spreading the same precursor solution on the surface of a substrate and hydrogel film. Detailed description of the synthesis and characterization of the adhesive hydrogels is given in the methods section. Notably, for the treatment of TM, a hydrogel patch must hold a trachea superficially and avoid collapsing as indicated by the numerical study. This essentially demands a hydrogel patch to possess excellent adhesive properties. To find the most suited adhesive hydrogel, we employed gelatin-coated glass slides (GCGS) to screen the formulation having the highest shear adhesion strength blend between HEAam and PEGDMA. Firstly, HEAam concentration was changed from 20 wt% to 50 wt% keeping covalent crosslinking density constant, as shown in Fig. 4b. We found that shear adhesion strength is increased ~4-fold from 80 kPa to more than 300 kPa after increasing the monomer concentration to 40 wt%. No improvement was seen afterwards. Therefore, 40 wt% HEAam concentration was selected to investigate the effect of crosslinker (PEGDM, 20kDa), as shown in Fig. 4c. We observed that adhesion strength on GCGS is enhanced from 180 kPa to more than 300 kPa by increasing PEGDM concentration from 0.25 wt% to 1.5 wt%, respectively. At 2.0 wt%, we observed no change in adhesion, however, it declined afterwards.

**Figure 4:**
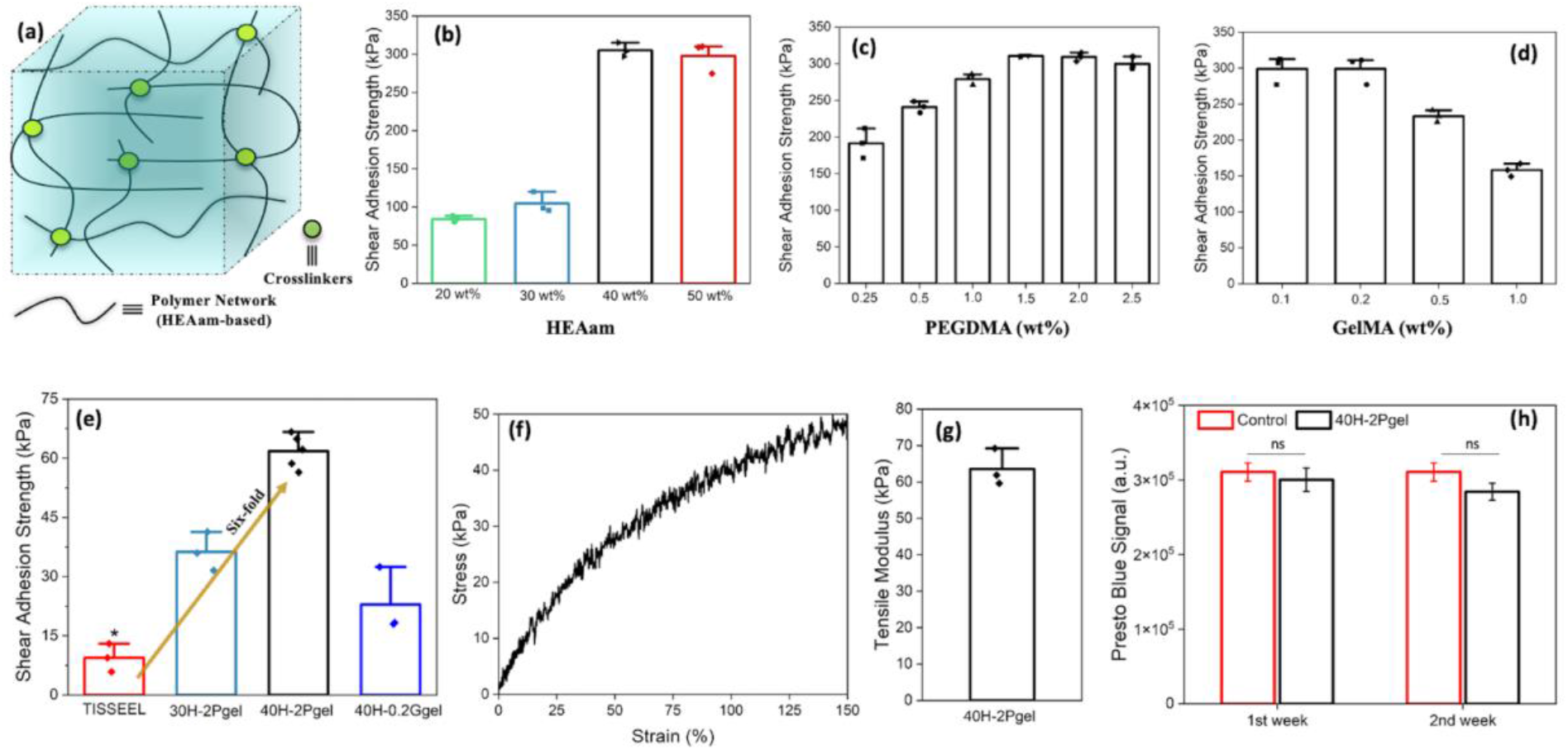
**(a)** Schematic of an adhesive hydrogel, **(b-d)** Screening to find out the hydrogel formulation having highest shear adhesion strength on the surface of gelatin-coated glass slides (GCGS), **(b)** by changing HEAam concentration at fixed PEGDMA concentration (2 wt%), **(c)** PEGDMA concentration at constant HEAam concentration (40 wt%), and **(d)** another crosslinker GelMA at constant HEAam concentration (40 wt%), **(e)** Shear adhesion strength of three hydrogels on the trachea and comparison with a commercial glue, TISSEEL. Undoubtedly, 40H-2Pgel hydrogel displayed best adhesion strength on the tracheal surface, (**f-g)** Typical stress-strain curve and tensile modulus of 40H-2Pgel hydrogel, **(h)** Cytotoxicity analysis of 40H-2Pgel after two weeks of incubation. *p<0.05 compared to other samples.

Hence, we selected 2 wt% PEGDMA as a final crosslinker concentration along with 40 wt% of HEAam to offset adhesion and bulk properties of our hydrogel called 40H-2Pgel. Likewise, we also investigated the effect of second crosslinker *i.e*., GelMA (0.1-1.0 wt%) keeping HEAam amount fixed at 40 wt% (Fig. 4d). We found that lower amount (0.1-0.2 wt%) of GelMA provides adhesion strength as strong as PEGDM (2 wt%) onto GCGS. Further addition of GelMA decreases the adhesion strength drastically. Therefore, 0.2 wt% GelMA along with 40 wt% of HEAam was chosen as second hydrogel called 40H-0.2Ggel. Subsequently, we re-examined the shear adhesion strength of these formulations on a wet rabbit-trachea surface. Fig. 4e shows the adhesion strength of each formulation that confirms the similar trend of adhesion and proved that 40H-2Pgel hydrogel has the highest shear strength of ~60 KPa on the wet tracheal surface (supplementary video 1). We also compared it against a commercial glue TISSEEL, a frequently used adhesive in surgery, and observed that 40H-2Pgel is a 6-fold stronger adhesive than TISSEEL on the tracheal surface, as shown in Fig. 4e.

The intrinsic adhesive properties of a hydrogel are due to the functional groups present on the polymer networks and their mechanical properties^[21–23]^. HEAam-based hydrogels (*i.e*., 40H-2Pgel) have two main functional groups -OH (hydroxyl) and -CONH-(amide) present on the polymer network^[24,25]^. These groups are crucial in forming multiple hydrogen bonding and strong van der Waals interactions on a trachea surface that also have numerous functional entities including primary amines (from lysine), carboxylic acids (from glutamic acid), thiols (from cysteine), and imidazole (from histidine)^[26,27]^. With high polymer (monomers) concentrations, a hydrogel should interact stronger with the tissue surface. This is why 40H-2Pgel showed higher adhesion on the tracheal surface than 30H-2Pgel. However, after reaching to an optimum amount of polymer concentration (40 wt% HEAam), the physical interactions between the two surfaces saturated and achieve a state where no further interactions can take place. On the other hand, covalent crosslinking dictates the bulk and dissipative (mechanical) properties of a hydrogel, which is a crucial aspect of achieving cohesive adhesion^[28,29]^. To achieve a robust and durable adhesion, it is equally important to balance out the physical interaction (due to surface functional groups present on the polymer chains of hydrogel) as well as cohesive interaction between hydrogels and tissues^[26,30]^. However, a high crosslink density decreases polymer chain length and associated chain mobility and forms a brittle hydrogel. Brittleness eventually reduces the hydrogel’s adhesion strength^[31]^. Therefore, we did not observe any adhesion improvement in hydrogel with high PEGDMA (> 2wt%) concentrations. GelMA, on the other hand, is a non-linear bulky protein that can crosslink a hydrogel network at multiple sites (not like PEGDMA). We believe that a high amount of GelMA will also lead to high crosslinking density to produce a brittle hydrogel^[32]^. Therefore, we selected 40H-2Pgel hydrogel to continue the further studies.

We examined the tensile properties of 40H-2Pgel hydrogel. The elastic modulus of hydrogel is ~65 kPa, as shown in Fig. 4f-g. We then sought to investigate the cytotoxicity of the hydrogel. Viability tests of mouse embryonic fibroblast cells were performed up to two weeks. We found that after two-weeks of incubation, hydrogel did not exhibit any cytotoxicity and showing statistically no difference in Presto blue signal compared to a control, as shown in Fig. 4h. This confirms the biocompatibility of 40H-2Pgel hydrogel, which is promising for the treatment of TM.

To confirm the numerical study and examine the potential use of hydrogel (possibly) to treat TM, we scanned the malacic trachea obtained after the enzymatic degradation (see methods for the details) with and without the application of the adhesive hydrogel under −15 mmHg pressure in μ-CT and compared the experimental results with the numerical model. The enzymatic degradation should mimic a mild malacic condition in trachea after softening its cartilage rings. Fig. 5a-b represents the synthesis and wrapping protocol of 40H-2Pgel hydrogel patch on a rabbit trachea for μ-CT measurements. To this end, a hydrogel splint (15×25×0.7 mm^3^) was obtained and wrapped around a trachea by performing a two-steps photo-polymerization. The two open ends of a trachea lumen were airtight using stoppers and zip-closure, as shown in Fig. 5b. Comparison between numerical and experimental results are shown in Fig. 5c-d. It can be observed that the overall shapes of the cross-section areas agree well between the μ-CT scans and the numerical results (Fig. 3e-f). The area of reconstructed and undeformed 11^th^ and 29.5^th^ tracheal rings was 29.5 and 24.3 mm^2^, respectively, which is similar to the values obtained from the numerical model (29.1 mm^2^ and 24 mm^2^). Under the same negative pressure (deformed state), the cross-sectional area of these rings decreased to 19.5 and 15 mm^2^, respectively. Likewise, the application of the adhesive hydrogel patch restrained the membrane movement and collapse by maintaining the airway area, specifically the inner area of the 11^th^ (23.6 mm^2^) and 29^th^ (20.4 mm^2^) rings is improved by 14% and 23%, respectively. These results validate the initial hypothesis and the numerical results. It is worth mentioning that even though the stiffness of 40H-2Pgel (~65 kPa, as shown in Fig. 4f-g) is one order of magnitude lower than that of the trachea cartilage, the adhesive property of hydrogel is crucial to limit the suction of the muscle membrane in the tracheal lumen. The pronounced results of the hydrogel patch assured that low-stiffness biocompatible materials with adhesive properties can be applied on the outer part of the membrane resulting in a considerable effect on restricting muscle deformation.

**Figure 5:**
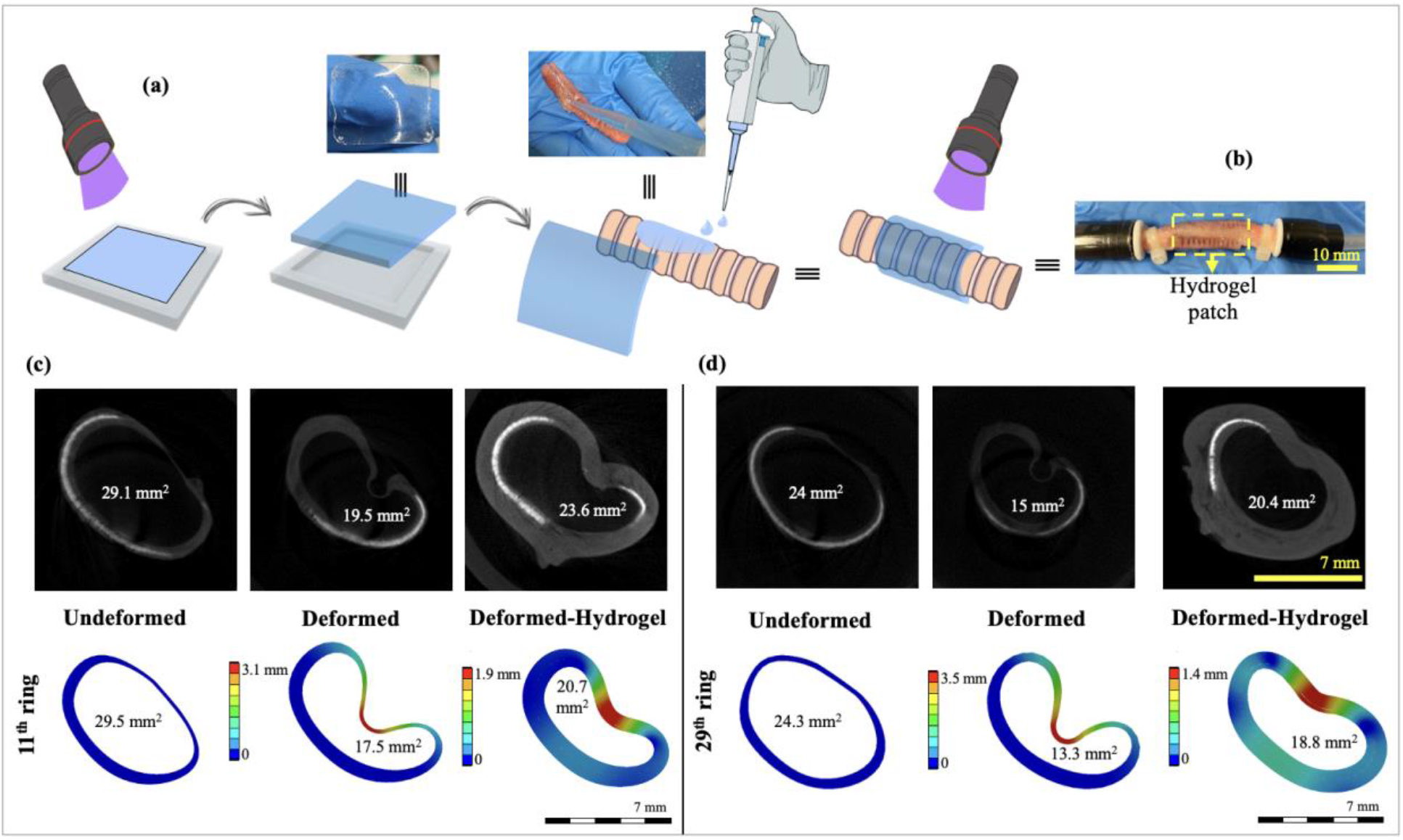
**(a)** Schematic of a hydrogel-patch (15×25×0.7 mm^3^) synthesis and preparation for a hydrogel-wrapped rabbit trachea, notably photo-polymerization was performed twice. For μ-CT analysis, rabbit trachea was kept in collagenase type 1 enzyme for 30 h to mimic mild TM conditions. **(b)** A photo of a hydrogel patch-wrapped trachea where both openings were fixed using plastic clips to create negative pressure. **(c-d)** μ-CT scans and numerical model of the 11^th^ and 29^th^ rings of trachea with and without application of the adhesive hydrogel at −15 mmHg. Results fall in line with our hypothesis that adhesive hydrogel patch has potential to correct trachea geometry and improve airway collapse.

The use of 40H-2Pgel hydrogel was further tested in *ex vivo* studies with rabbit tracheas. At this stage, we considered creating extreme malacic conditions (>75% collapse^[4]^) by completely removing eight to ten cartilage rings from healthy tracheas (harvested from a government-notified slaughterhouse) by the surgeons at the Lausanne University Hospital CHUV, Switzerland, (Fig. 6a-b). Subsequently, a negative pressure equivalent to the maximum physiological value (~5 kPa) was applied in a malacic trachea using a suction machine with an inbuilt manometer (Medela Surgicals, Switzerland) (Fig. 6c), and airway collapsing behavior was recorded using a flexible bronchoscope (Exalt, 6mm, BUN20EXALTBSRT100, Boston Scientific), with and without wrapping a hydrogel patch (15×35×0.7 mm^3^), as presented in Fig. 6d-e and 6g-h. Pictures of a trachea from the outside indicate that once a hydrogel patch is wrapped on a malacic trachea, the segment collapse was significantly lesser than without the hydrogel patch and comparable with the adjacent trachea (Fig. 6h).

**Figure 6:**
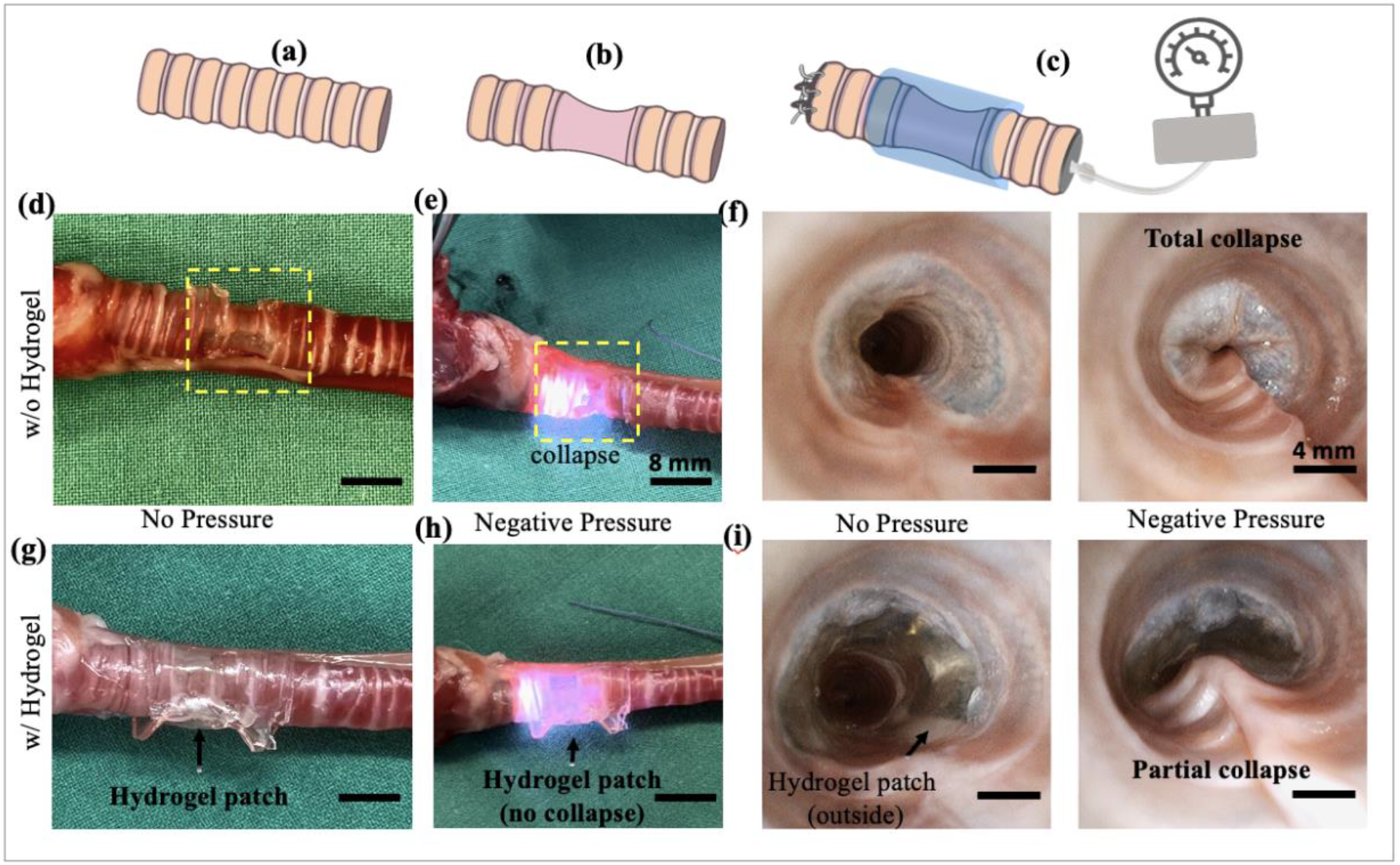
*Ex vivo* experiments: **(a)** Schematic of a normal trachea, **(b)** malacic trachea (tracheomalacia), **(c)** flexible bronchoscope with an in-built suction channel with the hydrogel patch wrapping the malacic trachea. The distal end of the trachea was closed with surgical sutures to allow maximal negative pressure effect on applying the suction. **(d)** An external view of a malacic trachea, cartilage rings were removed with preservation of the underlying mucosa to mimic tracheomalacia in a rabbit trachea, **(e)** an external image of a malacic trachea under pressure that immediately collapsed even at a very low negative pressure (−1 to −2 kPa). The bright light is of the bronchoscope that visualized the tracheal lumen (see supplementary video 2). **(f)** Endoluminal image of the trachea under negative pressure captured by bronchoscope (see supplementary video 3). Evidently, tracheal luminal structure was completely collapsed, a perfect example of the induced tracheomalacia, **(g)** An external image of hydrogel patch (15×35×0.7 mm^3^) wrapping the malacic trachea without any pressure, **(h)** An external view of the hydrogel patch-wrapped malacic trachea with negative pressure. The bright light of the bronchoscope can be seen through the transparent tracheal mucosa. Airway collapse improved up to 50%, (see supplementary video 4) and **(i)** endoscopic luminal images further confirmed the improvement in collapse (see supplementary video 5). Scale bars for external view of a malacic trachea and bronchoscope images with and without hydrogel are 8 and 4 mm, respectively.

We further recorded the inner luminal movement by the bronchoscope and observed the behavior of a malacic trachea under a negative pressure, as shown in Fig. 6f and 6i. Notably, without a hydrogel patch, airway was completely collapsed (Fig. 6f, supplementary videos 2 (outside) and 3 (inside)) whereas hydrogel wrapped malacic trachea shows up to 50% improvement in terms of airway opening (area = 82.5 mm^2^) compared to the adjacent non-operated trachea (area=170 mm^2^, fig. 6i, supplementary video 4 (outside) and 5 (inside)). This strongly suggests that the collapsing of a malacic trachea can be mechanically corrected by wrapping an adhesive hydrogel patch extraluminally.

## Conclusion

Tracheomalacia (TM) is indeed a life-threatening situation for newborns. Established treatment methods come with various flaws; thus, a better and more realistic approach is paramount. Herein, we proposed the use of adhesive hydrogel patch to prevent airway collapse by supporting a malacic trachea externally. Initially, the numerical study pointed out that the application of an adhesive hydrogel patch can help to maintain physiological shape of a malacic trachea and to constrain the membrane during collapse. Based on numerical findings, new adhesive hydrogels were formulated after employing the hydroxyethyl acrylamide (HEAam) and polyethylene glycol methacrylate (PEGMDA) as main polymer network and crosslinker, respectively. The shear adhesion of the biocompatible 40H-2Pgel hydrogel on tracheal surface was found to be more than 60 kPa. Once hydrogel patch is wrapped around a malacic trachea, micro-CT and *ex vivo* measurements confirmed that the airway collapsing can be improved and is a promising finding for the correction of TM in newborns.

Although this study opens up a new approach to correct TM, further investigation about the hydrogel properties needs to be addressed before going into *in vivo* studies. Since the trachea is in contact with the body fluid, adhesive hydrogel should not show excessive swelling or degradation in order to maintain its mechanical properties, especially adhesion. Moreover, the fatigue resistance of the adhesive hydrogel under dynamic tracheal movement will be investigated to observe the long-term performance.

## Acknowledgements

We would like to thank Meinrad Odermatt of Delimpex AG, Pfäffikon, Switzerland for providing the rabbit tracheas. Arnaud Bichat and the animal experimentation platform of the CHUV, Lausanne for their help in *ex vivo* experiments. This study is supported by a Sinergia grant from the Swiss National Science Foundation (#CRSII5_189913 /1).

---

**Supporting video 1:** Shows the robust adhesion of 40H-2Pgel hydrogel patch on a wet rabbit trachea, ready for the micro-CT scan.

**Supporting video 2:** Shows the extreme malacic condition created in a rabbit trachea after removing the cartilage rings and demonstrates the collapsing behavior under the negative pressure (−5 kPa).

**Supporting video 3:** Shows the endoluminal video of the same malacic trachea (used in video 2) recorded by the bronchoscope. The recording confirms the complete luminal collapse under the negative pressure (−5 kPa).

**Supporting video 4:** Shows a patch of 40H-2Pgel hydrogel is wrapped around the same malacic trachea (used in video 2 & 3). Partial collapse could be seen when the same negative pressure is applied (−5 kPa).

**Supporting video 5**: Shows the endoluminal video of the 40H-2Pgel hydrogel wrapped around malacic trachea (used in video 2 & 3) recorded by the bronchoscope. Again, the recording confirms partial collapse under the same negative pressure (−5 kPa).

## Experimental and Methods Section

### Materials

*N*-(2-Hydroxyethyl)-acrylamide (HEAam) was purchased from Chemie Brunschweig AG. Lithium-Phenyl-2,4,6 trimethylbenzoylphosphinat (LAP) was obtained from Sigma Aldrich. Polyethyleneglycoldimethacrylate (PEGDMA, M_n_ = 20kDa) was purchased from Polysciences. Gelatin Type A from porcine skin (ref. G2500), methacrylic anhydride, sodium hydroxide, dialysis sacks (MWCO 6000– 8000 Da), and hydrochloric acid were purchased from Sigma Aldrich. Young rabbit tracheas (4-6 Kg) were provided by Delimpex AG, Pfäffikon, Switzerland

### Fabrication of Adhesive Hydrogel

HEAam (20-50% w/v), PEGDMA (0.25 to 2.5% w/v), and photoinitiator LAP (0.05% w/v) were dissolved in PBS using vortex in absence of light. Gelatin methacrylate (GelMA) was synthesized following the protocol reported earlier^[33]^. The hydrogel precursor was poured into a 15×15×0.7 mm^3^ custom-made Teflon molds and covered with plastic slides. Polymerization was achieved by the illumination of 405 nm light (3 mW cm^-2^) for 2 minutes.

### Adhesion Measurements

Shear adhesion set-up based on ASTM F2255 standards^[34]^ were used to measure shear adhesion of the hydrogels on gelatin-coated glass slides (GCGS 10×25 mm^2^) and rabbit trachea.

#### Glass adhesion

The preformed hydrogel patch of 15×15×0.7 mm^3^ was obtained as described in fabrication section. 100 μL of same precursor solution was poured on the preformed hydrogel surface already adhered to a GCGS and second GCGS was fixed on the hydrogel surface followed by the second photo-polymerization for 2 minutes. After second polymerization, adhesion measurements were performed using 50 N load cell connected to Instron E3000 mechanical testing machine with constant loading rate of 1 mm/s. Shear adhesion strength of the hydrogels was calculated dividing the maximum load by the surface area of the hydrogel patch.

#### Adhesion on Trachea

Rabbit tracheas were hydrated into PBS for 45 minutes before the adhesion measurements and cut into the dimension of 10×25 mm^2^ (Fig. S1a). Rabbit trachea piece was glued on the glass slide using Superglue (Loctite 401, Fig. S1b). Once a hydrogel patch (15×15×0.7 mm^3^) is prepared (as mentioned earlier), 100 μL of the precursor solution was poured onto wet trachea and hydrogel patch was placed on tissue surface avoiding any air bubble formation (Fig. S1c). Furthermore, 100 μL of the precursor solution was poured again on the top surface of the hydrogel patch and covered by glass slide (Fig.S1d) followed by second photo-polymerization step for 2 minutes (Fig. S1d).

**Figure S1:**
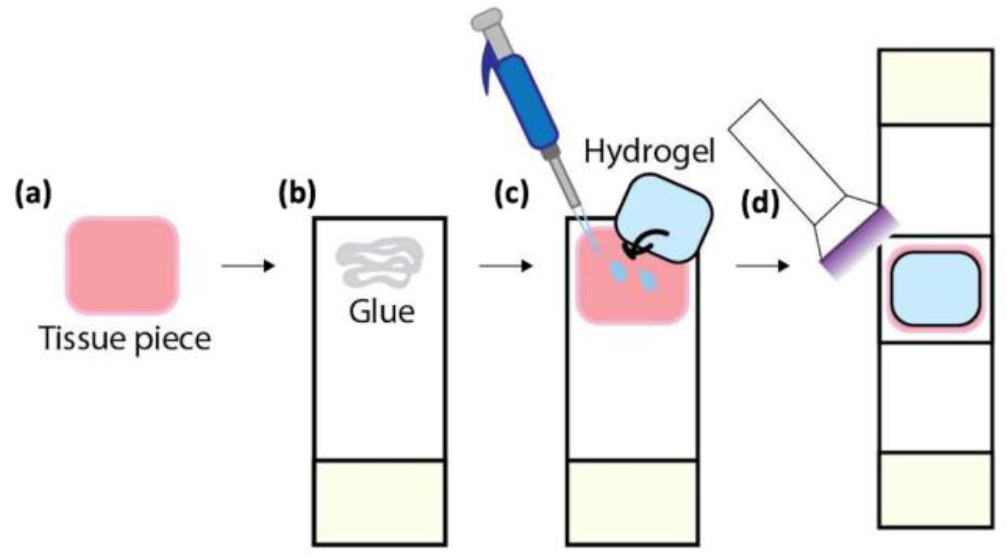
(a-b) Trachea pieces were glued on glass slide, (c) 100 μL of precursor was poured onto tissue surface, hydrogel was placed on it and 100 μL of precursor was poured again on top surface of the hydrogel patch, (d) glass slide was put onto hydrogel and second polymerization was followed.

### Numerical Study

#### Geometry

The geometry of the trachea was derived from one of the healthy rabbit tracheas (4-6 Kg) that were scanned in the experimental study and was modelled on AutoCAD Inventor. In order to accurately capture the geometrical features of the trachea, like the radially variable thickness and the general asymmetric shape, five equidistant scans were used as a guideline to sketch the exact profile at the corresponding cross-sections. A 3D model was then fitted through these sketches using a loft function. Subsequently, the 3D volume was divided in 44 regions (corresponding to 22 cartilage rings, 21 connective tissue and 1 muscle domains). To maintain the same ratio of cartilage-to-tissue volume, the partitioning process was based on an independent 3D reconstruction of the tracheal cartilage rings, which was produced on the open-source software Slicer.

#### Boundary and loading conditions

To facilitate the correlation between the experimental and the numerical results, the imposed boundary conditions on both ends of the trachea were chosen to be fixed. Thus, the longitudinal deformations were eliminated, but most importantly, any unwanted radial translations of the trachea which would be caused by the naturally present asymmetries, were minimized. Finally, the pressure load was applied on the internal surface of the geometry, facing the inward direction in order to describe the systolic behaviour during expiration. Based on the literature, the pressure experienced by the inner trachea walls during expiration is around −15 mmHg, hence it was the pressure of interest for the present numerical study.

#### Material properties

The Mooney-Rivlin model was selected for the present study as mentioned in the manuscript while the parameters inside Ansys were tuned according to the uniaxial stress-strain curves of a human trachea^[20]^ and were assigned to the corresponding parts (Table S1). These parameters vary depending on the animal, age and gender^[35]^ and could be further optimized for subject-specific measurements, but such a procedure lies beyond the scope of the present study.

**Table S1:** Fitted Mooney-Rivlin parameters, following the uniaxial stress-strain curves of C_ij_ are material constants and D1 is the incompressibility parameter, as defined in the Ansys Mechanical solver.

|  | Cartilage | Muscle | Tissue |
| --- | --- | --- | --- |
| $C_{10}$ (MPa) | 0.440885 | 0.10502727 | 1.367453 |
| $C_{01}$ (MPa) | 1.894736 | -0.0180258 | -1.14558 |
| $C_{11}$ (MPa) | 0.342602 | 0.05891425 | -0.09921 |
| $D_1$ (MPa<sup>-1</sup>) | 0 | 0 | 0 |

#### Computational mesh

Through experimentation, the mesh strategy was proven to be an important factor for the stability but also for the convergence of the solution process. Several different strategies were attempted, including dynamic remeshing, and the most robust of which was to divide the geometry in the longitudinal domains of variable mesh density, according to their expected relative deformation, as shown in Fig. S2a-b. Hexahedral elements were used where possible, due to their higher accuracy compared to tetrahedrons, while the quadratic element order was preferred to the linear, due to its ability to capture element bending^[36]^.

**Figure S2:**
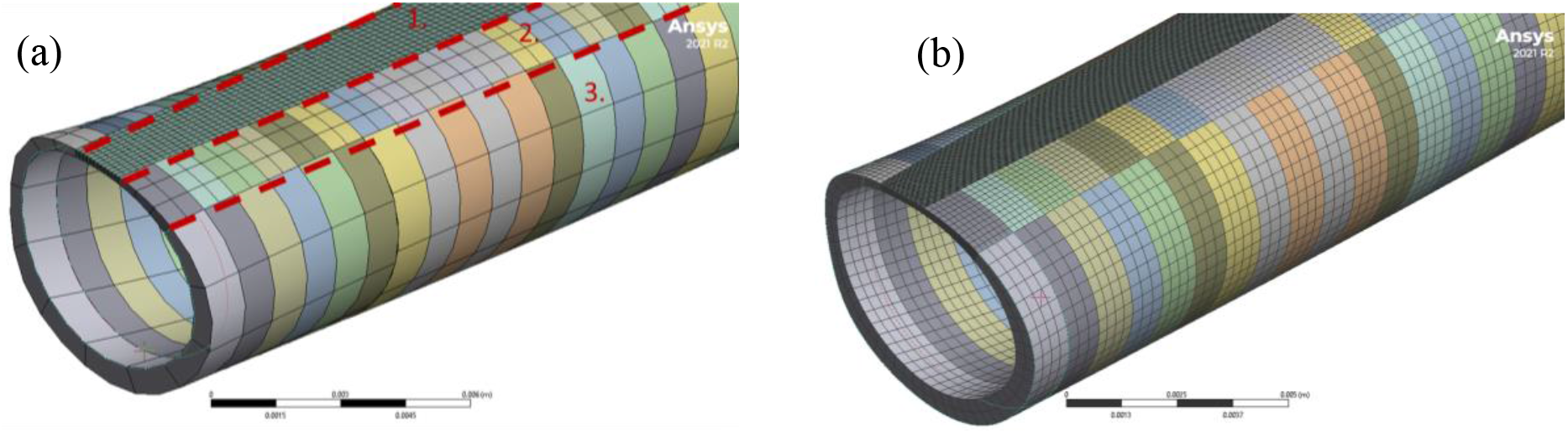
Coarse (a) and fine (b) computational meshes of the trachea and their corresponding refinement regions (1-3 from low to high element size).

**Figure S3:**
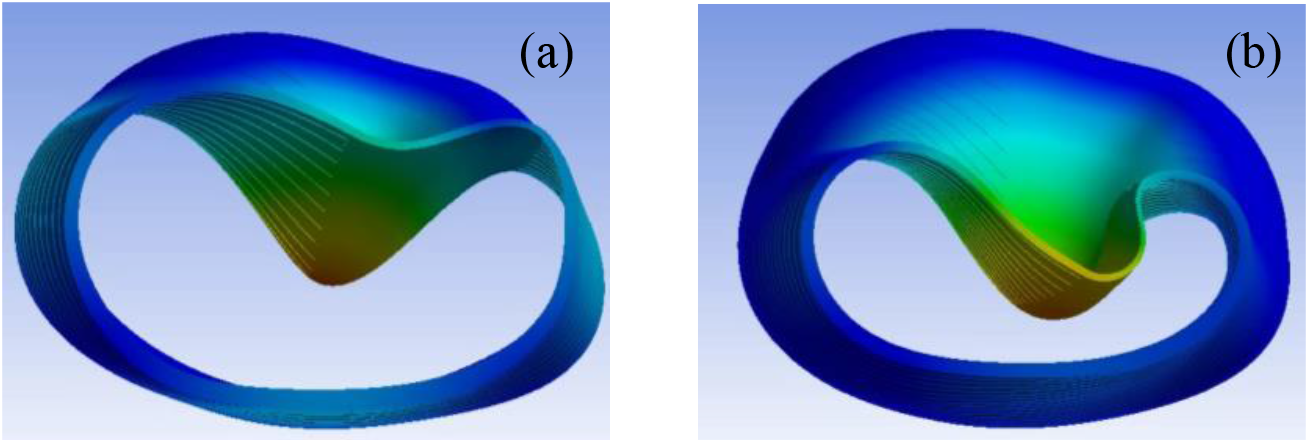
Comparison of the behaviour of a linear (a) vs a large deformation model (b). Since the deformations are comparable to the diameter of the trachea, the linear model does not capture the features of the deformation as observed in the experiments.

#### Deformation model

For the simulations, a static solver was selected since the main focus is aimed on the static behavior of the trachea under peak loading. However, the expected deformations are comparable to the radial direction (Fig. S3a-b), which is the main direction of interest for malacic tracheas. This induces further geometrical non-linearities on the system, which require an iterative calculation of the stiffness matrix *K* at each pseudo-time step. For this reason, the large deflection model is selected in the software settings. It is worth mentioning that although using a linear model might converge to a rational solution, the internal strain energy of the material is not conserved. The comparison between a linear and a non-linear solution is shown in (Supplementary Fig. S3a-b), where it becomes apparent that a linear deformation model cannot accurately capture the shrinking of the trachea or the membrane folding (which were both observed in the experiments), while the non-linear model achieves both. For each sub-step of the solution, a direct solver is selected, as it has a positive impact on the computational time performance. The appropriate dynamic time-step for each pseudo-steady iteration is within the range of 10^-5^ and 4·10^-3^ sec and it is chosen automatically by the solver.

#### Mesh independence

To achieve mesh-independent results, a parametric study of multiple refinement steps was conducted, where the average element size on each individual meshing domain was multiplied by a factor of 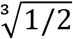, in order to obtain a constant doubling of the total element number at each refinement. As shown in Fig. S4a, both the maximum deformation and the maximum von Mises stress converges after about 2·10^5^ elements. It can also be observed that the percentage of deformation improvement between the coarsest and the finest meshes is not higher than 2.5% (Fig. S4b), which is an indication for the good quality of the mesh, as well as the suitability of the adopted domain division strategy. The runs were performed on a Ryzen 9 5900HX, while the timings for the coarsest and the finest runs were 3 min and 16 hours, respectively. Thus, an intermediate model of 160·10^3^ elements was selected which took around 6 hours to be solved. It should be mentioned that for parametric runs regarding the shape of the trachea, a coarser mesh could be used since the relative variation of the membrane deformation appears to be small (Fig. S4b).

**Figure S4:**
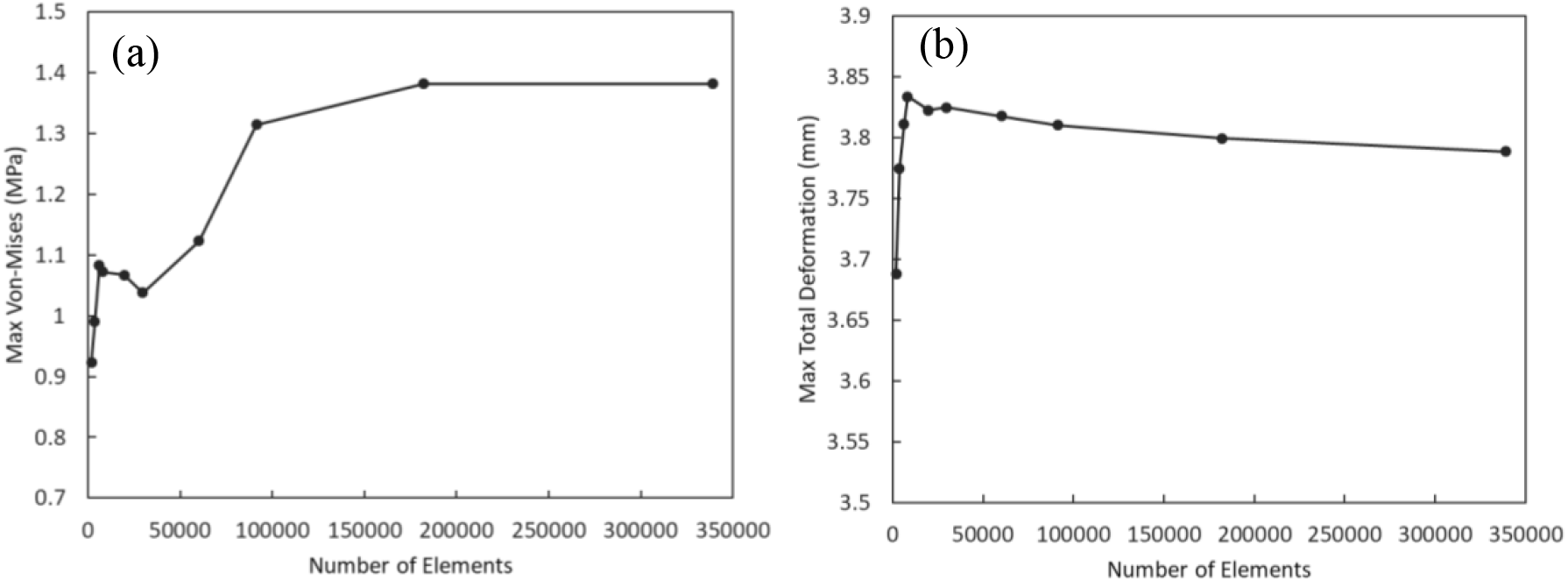
Convergence of maximum von Mises stress (a) and maximum total deformation (b) due to the gradual mesh refinement. A density of 160·10^3^ elements produce mesh-independent results.

#### Micro (μ)-CT Measurements

Two ends of the trachea were fixed to the stoppers and tightened with zip closure to make the trachea airtight. After that, healthy, malacic trachea and/or trachea with and without hydrogel patch (15×25×0.7 mm^3^) were placed inside the μ-CT machine (Skyscan). The syringe was connected to the trachea to apply positive and negative pressure, and an electronic manometer was also connected to the plastic tubes to read the applied pressure values. Open area of the airway was calculated using ImageJ 1.51 software (National Institute of Health, Bethesda, Maryland).

#### Tensile test

The tensile tests of dog-bone shaped hydrogel specimens (2 mm thickness, 5 mm neck width and 4.5 mm gauge length) were carried out using an Instron E3000 linear testing machine (Norwood, MA, USA) with a 50 N load cell. The specimens were placed within the grippers and elongated at a loading rate of 0.1 mm. s^-1^. The elastic modulus (tensile modulus) of the hydrogels was calculated on the initial linear slope of the tensile stress-strain curves at 10–30% strain (n = 4).

#### Enzymatic Degradation of Tracheal Cartilage

To mimic the mild TM condition in a healthy rabbit trachea for μ-CT measurements, tracheal cartilages were degraded using 0.1% (w/v) collagenase-I (col-I) solution. Briefly, 500 μL of col-I solution was poured into a petri dish and fresh rabbit trachea was placed in it facing tracheal membrane upwards without interacting with col-I solution. To avoid dehydration, a custom-made water bath was used to float a petri dish (having trachea) on the water during degradation process. Degradation process was conducted for 12, 18, 24 and 48 hours. After each time interval, trachea was removed from the solution and washed thoroughly with PBS. Degraded (malacic) tracheas were stored at −20°C before further use.

#### Ex vivo Experiments

To mimic the extreme TM condition, 8 to 10 cartilage rings were removed from a frozen and then thawed rabbit tracheas with scalpel without damaging the inner tracheal mucosa. The distal end of the trachea was closed using surgical sutures and ensure an air-tight condition. The flexible bronchoscope was introduced into the trachea through the laryngeal inlet. Hydrogel patch (15×35×0.7 mm^3^) was prepared as described earlier. Then, precursor solution was poured on the trachea surface and wrapped with the hydrogel patch avoiding any air bubble followed by the second photo-polymerization using a portable torch for 2 minutes. Then, a negative pressure (−5 kPa, maximum physiological pressure) was applied to the trachea by suction set-up (Medela Surgicals) and collapse behavior was observed and recorded by a flexible bronchoscope (Boston Scientific). Experiment was repeated 3 times applying 20 +/- pressure cycles. Open area of the airway was calculated using ImageJ 1.51 software (National Institute of Health, Bethesda, Maryland).

#### Toxicity Test

Mouse embryonic fibroblast cells (NiH3T3, passage number 4) were used to evaluate biocompatibility of the adhesive hydrogel. Briefly, the precursor of the adhesive hydrogel was filtered inside the laminar flow and poured into the sterilized disk-shaped molds (∅ 5mm). Then, molds were covered with glass slides and polymerization was performed for 2 minutes inside the laminar flow. After polymerization, the adhesive hydrogels were placed into 24-well plate filled with complete cell culture medium (DMEM supplemented with 10% (v/v) Fetal Bovine Serum, 1% (v/v) Penicillin Streptomycin, 1% (v/v) L-Glutamine). Subsequently, the cell culture plate was placed into the incubator (37 °C and 5% CO_2_) for 1 and 2 weeks. After fixed time intervals, hydrogels were removed from the plate. The conditioned medium was put into 96 well-plates containing 1000 cells/well and incubated again for 1 day. After day one, medium was aspirated from the well-plate and 100 μL of 10% (v/v) PrestoBlue (A13261, Life Technologies) was put into each well and the plate was incubated for 30 min. After that, fluorescence was measured at 595 nm using a microplate reader (Wallac 1420 Victor2, PerkinElmer). Toxicity experiments were performed in triplicates using five replicates for each experiment. DMEM solution with 1000 cells/well was taken as control.

#### Statistical Analysis

The OriginLab software (Northampton, MA) was used for statistical analysis of the data. One-way analysis of variance (ANOVA) with Tukey’s test was applied for data analysis. p < 0.05 were considered statistically significant.

## References

[1] C. Serrano-Casorran, S. Lopez-Minguez, S. Rodriguez-Zapater, C. Bonastre, J. A. Guirola, M. A. De Gregorio, Pediatr. Pulmonol. 2020, 55, 1757.

[2] A. Kamran, R. W. Jennings, Front. Pediatr. 2019, 7, 1.

[3] S. Choi, C. Lawlor, R. Rahbar, R. Jennings, JAMA Otolaryngol. - Head Neck Surg. 2019, 145, 265.

[4] G. Burg, M. M. Hossain, R. Wood, E. B. Hysinger, Ann. Am. Thorac. Soc. 2021, 18, 1749.

[5] S. D. Murgu, H. G. Colt, Respirology 2006, 11, 388.

[6] E. B. Hysinger, Curr. Probl. Pediatr. Adolesc. Health Care 2018, 48, 113.

[7] W. Svetanoff, R. Jennings, J. Lung Heal. Dis. 2018, 2, 17.

[8] A. Kamran, C. J. Smithers, C. W. Baird, R. W. Jennings, JTCVS Tech. 2021, 8, 160.

[9] C. Wallis, E. Alexopoulou, J. L. Antón-Pacheco, J. M. Bhatt, A. Bush, A. B. Chang, A. M. Charatsi, C. Coleman, J. Depiazzi, K. Douros, E. Eber, M. Everard, A. Kantar, I. B. Masters, F. Midulla, R. Nenna, D. Roebuck, D. Snijders, K. Priftis, Eur. Respir. J. 2019, 54, 1.

[10] F. Gorostidi, A. Reinhard, P. Monnier, K. Sandu, Laryngoscope 2016, 126, 2605.

[11] F. Gorostidi, C. Courbon, M. Burki, A. Reinhard, K. Sandu, Laryngoscope 2018, 128, E53.

[12] X. Zhang, G. Pan, Ocean Eng. 2020, 213, DOI 10.1016/j.oceaneng.2020.107705.

[13] T. Christo Michael, A. R. Veerappan, S. Shanmugam, Eng. Fract. Mech. 2012, 79, 138.

[14] R. Ramasamy, T. M. Y. S. Tuan Ya, MATEC Web Conf. 2014, 13, DOI 10.1051/matecconf/20141302003.

[15] S. Rafieian, H. Mirzadeh, H. Mahdavi, M. E. Masoumi, IEEE J. Sel. Top. Quantum Electron. 2019, 26, 154.

[16] Q. Chen, H. Chen, L. Zhu, J. Zheng, J. Mater. Chem. B 2015, 3, 3654.

[17] V. K. Rana, A. Tabet, J. A. Vigil, C. J. Balzer, A. Narkevicius, J. Finlay, C. Hallou, D. H. Rowitch, H. Bulstrode, O. A. Scherman, ACS Macro Lett. 2019, 8, 1629.

[18] P. Bagnoli, F. Acocella, M. Di Giancamillo, R. Fumero, M. L. Costantino, J. Biomech. 2013, 46, 462.

[19] P. A. L. S. Martins, R. M. N. Jorge, A. J. M. Ferreira, Strain 2006, 42, 135.

[20] F. Safshekan, M. Tafazzoli-Shadpour, M. Abdouss, M. B. Shadmehr, F. Ghorbani, Int. J. Solids Struct. 2020, 190, 35.

[21] P. Karami, N. Nasrollahzadeh, C. Wyss, A. O’Sullivan, M. Broome, P. Procter, P. E. Bourban, C. Moser, D. P. Pioletti, Macromol. Rapid Commun. 2021, 42, DOI 10.1002/marc.202000660.

[22] J. Li, A. D. Celiz, J. Yang, Q. Yang, I. Wamala, W. Whyte, B. R. Seo, N. V. Vasilyev, J. J. Vlassak, Z. Suo, D. J. Mooney, Science (80-.). 2017, 357, 378.

[23] H. Yuk, C. E. Varela, C. S. Nabzdyk, X. Mao, R. F. Padera, E. T. Roche, X. Zhao, Nature 2019, 575, 169.

[24] X. Yu, Y. Li, J. Yang, F. Chen, Z. Tang, L. Zhu, G. Qin, Y. Dai, Q. Chen, Macromol. Mater. Eng. 2018, 303, 1.

[25] X. Sun, S. He, M. Yao, X. Wu, H. Zhang, F. Yao, J. Li, J. Mater. Chem. C 2021, 9, 1880.

[26] S. Nam, D. Mooney, Chem. Rev. 2021, DOI 10.1021/acs.chemrev.0c00798.

[27] V. G. Muir, J. A. Burdick, Chem. Rev. 2021, DOI 10.1021/acs.chemrev.0c00923.

[28] M. B. Yi, T. H. Lee, G. Y. Han, H. Kim, H. J. Kim, Y. Kim, H. S. Ryou, D. U. Jin, ACS Appl. Polym. Mater. 2021, 3, 2678.

[29] P. Karami, C. S. Wyss, A. Khoushabi, A. Schmocker, M. Broome, C. Moser, P. E. Bourban, D. P. Pioletti, ACS Appl. Mater. Interfaces 2018, 10, 38692.

[30] J. Shen, H. Zhang, J. Zhu, Y. Ma, H. He, F. Zhu, L. Jia, Q. Zheng, Front. Chem. 2022, 10, 1.

[31] J. H. Lee, T. H. Lee, K. S. Shim, J. W. Park, H. J. Kim, Y. Kim, S. Jung, Int. J. Adhes. Adhes. 2017, 74, 137.

[32] A. Mathur, A. Li, V. Maheshwari, J. Phys. Chem. Lett. 2021, 12, 1481.

[33] M. Zhu, Y. Wang, G. Ferracci, J. Zheng, N. J. Cho, B. H. Lee, Sci. Rep. 2019, 9, 1.

[34] ASTM International, https://doi.org/10.1520/F2255-05R15 (accessed: April 2021)

[35] C. Zweben, P. Beaumont, Comprehensive Composite Materials II, Elsevier, (2017).

[36] F. Safshekan, M. Tafazzoli-Shadpour, M. Abdouss, M. B. Shadmehr, Materials (Basel). 2016, 9.

